# Determinants of Real-Time fMRI Neurofeedback Performance and Improvement – a Machine Learning Mega-Analysis

**DOI:** 10.1101/2020.10.21.349118

**Authors:** Amelie Haugg, Fabian M. Renz, Andrew A. Nicholson, Cindy Lor, Sebastian J. Götzendorfer, Ronald Sladky, Stavros Skouras, Amalia McDonald, Cameron Craddock, Lydia Hellrung, Matthias Kirschner, Marcus Herdener, Yury Koush, Marina Papoutsi, Jackob Keynan, Talma Hendler, Kathrin Cohen Kadosh, Catharina Zich, Simon H. Kohl, Manfred Hallschmid, Jeff MacInnes, Alison Adcock, Kathryn Dickerson, Nan-Kuei Chen, Kymberly Young, Jerzy Bodurka, Michael Marxen, Shuxia Yao, Benjamin Becker, Tibor Auer, Renate Schweizer, Gustavo Pamplona, Ruth A. Lanius, Kirsten Emmert, Sven Haller, Dimitri Van De Ville, Dong-Youl Kim, Jong-Hwan Lee, Theo Marins, Megumi Fukuda, Bettina Sorger, Tabea Kamp, Sook-Lei Liew, Ralf Veit, Maartje Spetter, Nikolaus Weiskopf, Frank Scharnowski, David Steyrl

## Abstract

Real-time fMRI neurofeedback is an increasingly popular neuroimaging technique that allows an individual to gain control over his/her own brain signals, which can lead to improvements in behavior in healthy participants as well as to improvements of clinical symptoms in patient populations. However, a considerably large ratio of participants undergoing neurofeedback training do not learn to control their own brain signals and, consequently, do not benefit from neurofeedback interventions, which limits clinical efficacy of neurofeedback interventions. As neurofeedback success varies between studies and participants, it is important to identify factors that might influence neurofeedback success. Here, for the first time, we employed a big data machine learning approach to investigate the influence of 20 different design-specific (e.g. activity vs. connectivity feedback), region of interest-specific (e.g. cortical vs. subcortical) and subject-specific factors (e.g. age) on neurofeedback performance and improvement in 608 participants from 28 independent experiments.

With a classification accuracy of 60% (considerably different from chance level), we identified two factors that significantly influenced neurofeedback performance: Both the inclusion of a pre-training no-feedback run before neurofeedback training and neurofeedback training of patients as compared to healthy participants were associated with better neurofeedback performance. The positive effect of pre-training no-feedback runs on neurofeedback performance might be due to the familiarization of participants with the neurofeedback setup and the mental imagery task before neurofeedback training runs. Better performance of patients as compared to healthy participants might be driven by higher motivation of patients, higher ranges for the regulation of dysfunctional brain signals, or a more extensive piloting of clinical experimental paradigms. Due to the large heterogeneity of our dataset, these findings likely generalize across neurofeedback studies, thus providing guidance for designing more efficient neurofeedback studies specifically for improving clinical neurofeedback-based interventions. To facilitate the development of data-driven recommendations for specific design details and subpopulations the field would benefit from stronger engagement in Open Science and data sharing.

## Introduction

Real-time functional magnetic resonance (fMRI) neurofeedback is a non-invasive technique that enables healthy individuals and patients to voluntarily regulate neural signals. In the last decades, this method has gained growing popularity in the neuroimaging community and, to date, a wide range of real-time fMRI neurofeedback studies have collectively demonstrated the feasibility of volitional regulation through real-time fMRI neurofeedback (see Thibault, MacPherson, Lifshitz, Roth, & Raz (2018)). Further, many of these studies have also shown behavioral changes in healthy individuals, as well as clinical improvements in patient populations after neurofeedback training. In healthy participants, real-time fMRI neurofeedback training has been specifically linked to improvements in attention (e.g. DeBettencourt, Cohen, Lee, Norman, & Turk-Browne, 2015; Pamplona et al., 2020), emotion regulation (Koush et al., 2015; Paret & Hendler, 2020; Zich et al., 2020), memory (e.g. Scharnowski et al., 2015; Sherwood, Kane, Weisend, & Parker, 2016; Zhang, Yao, Zhang, Long, & Zhao, 2013), motivation (e.g. Zhi et al., 2018), motor performance (e.g. Bray, Shimojo, & O’Doherty, 2007; Scharnowski et al., 2015; Sitaram et al., 2012; Zhao et al., 2013), speech performance (Rota et al., 2009), and visual perception (e.g. Scharnowski, Hutton, Josephs, Weiskopf, & Rees, 2012; Shibata, Watanabe, Sasaki, & Kawato, 2011). In clinical populations, real-time fMRI neurofeedback training has been shown to both improve clinical measures and normalize pathological neural characteristics in patients suffering from a wide range of disorders, such as alcohol and nicotine addiction (Canterberry et al., 2013; Hanlon et al., 2013; Hartwell et al., 2016; Karch et al., 2015; Kim, Yoo, Tegethoff, Meinlschmidt, & Lee, 2015; X. Li et al., 2013), anxiety (Morgenroth et al., 2020), borderline personality disorder (Paret et al., 2016), depression (Linden et al., 2012; Quevedo et al., 2020; Young et al., 2017, 2014), obsessive compulsive disorder (Buyukturkoglu et al., 2015), phobia (Zilverstand, Sorger, Sarkheil, & Goebel, 2015), post-traumatic stress disorder (Gerin et al., 2016; Nicholson et al., 2017), schizophrenia (Bauer et al., 2020), obesity (Frank et al., 2012), chronic pain (deCharms et al., 2005; Guan et al., 2014), Huntington’s disease (Papoutsi et al., 2018), Parkinson’s disease (Buyukturkoglu et al., 2013; Subramanian et al., 2011), tinnitus (Emmert, Kopel, et al., 2017; Haller, Birbaumer, & Veit, 2010), and visuo-spatial neglect (Fabien Robineau et al., 2019).

Critically however, not all participants undergoing real-time fMRI neurofeedback training optimally benefit from the aforementioned improvements on behavioral and clinical measures, due to variations in their success on acquiring neural control. Previous real-time fMRI neurofeedback studies have reported relatively high rates of non-responders, i.e., participants who fail to regulate their brain signals in the desired direction (Bray et al., 2007; Chiew, LaConte, & Graham, 2012; deCharms et al., 2005; Johnson et al., 2012; Ramot, Grossman, Friedman, & Malach, 2016; F. Robineau et al., 2014; Scharnowski et al., 2012; Yoo, Lee, O’Leary, Panych, & Jolesz, 2008). Averaging across these studies, the non-responder rate of real-time fMRI neurofeedback studies is estimated to lie around 38% (Haugg et al., 2020). Here, it should be noted that, to date, no standard thresholds for identifying non-responders are available and definitions of non-responders often vary between studies. Generally, even realtime fMRI neurofeedback participants who were eventually able to gain control over their own brain signals still showed large variability in their neurofeedback regulation performance (Haugg et al., 2020). Similar estimations and observations have also been reported in the electroencephalogram (EEG) neurofeedback literature, where the so-called “neurofeedback inefficacy problem” refers to the variability in neurofeedback success and comprises a well-known issue (Alkoby, Abu-Rmileh, Shriki, & Todder, 2017). Therefore, the fields of both EEG- and fMRI-based neurofeedback would greatly benefit from methodologically advanced investigations that can reveal the factors responsible for the unexplained variability of neurofeedback success.

Interestingly, previous studies have demonstrated that the proportion of responders varies between different neurofeedback studies. Of importance, this suggests that some neurofeedback study-specific parameters might be more beneficial for neurofeedback success than others. Previously, few empirical studies have investigated the influence of neurofeedback design parameters on neurofeedback success. Specifically, two independent studies found that using an intermittent feedback display was superior over using a continuous feedback display (Hellrung et al., 2018; Johnson et al., 2012), while conversely, a third study reported this effect only for a single session of neurofeedback, but not for multiple neurofeedback sessions (Emmert, Kopel, et al., 2017). In another study, Papoutsi and colleagues investigated the influence of activity-versus connectivity-based neurofeedback on neurofeedback success, but did not find a significant difference between activity- and connectivity-based neurofeedback (Papoutsi et al., 2020). Interestingly, Kim et al. reported increased neurofeedback efficacy when combining connectivity-based information with activity-based neurofeedback (Kim et al., 2015). Focusing on subject-specific psychological factors in a systematic review, Cohen Kadosh and colleagues observed that attention and motivation might be important factors for determining neurofeedback success (Cohen Kadosh & Staunton, 2019). However, an empirical validation of these suggestions is still needed. Other empirical studies observed a relationship between subject-specific questionnaires and neurofeedback success, yet these questionnaires were highly specific for the trained target region and participant population, and therefore do not generalize to other neurofeedback studies (Emmert, Breimhorst, et al., 2017; Koush et al., 2015).

Taken together, these empirical studies contribute invaluable information regarding the optimal design of neurofeedback studies. However, many critical factors that might influence neurofeedback success have not been investigated yet. For instance, it is not known whether a large number of neurofeedback training runs is beneficial for neurofeedback success, an essential question in the field of fMRI-based neurofeedback due to the high cost of scanning hours. This also includes the question of whether neurofeedback training should be performed across several training days to facilitate neurofeedback learning through sleep consolidation. Other important factors are the inclusion of reinforcers such as monetary rewards (Sepulveda et al., 2016) and social rewards (Mathiak et al., 2010), or the highly debated question of whether participants should receive precise or more open instructions regarding regulation strategies (Sitaram et al., 2016). Ultimately, the number of possible factors that might influence neurofeedback performance and the number of conceivable interactions between these factors are immense and it would not be feasible to untangle them and optimize design empirically. Further, statistical power and generalizability across different study designs are limited in original empirical studies.

On balance, ‘big data’ approaches encompassing a wide range of neurofeedback participants and studies constitute an unprecedented opportunity that can be used to investigate a large number of factors that might influence neurofeedback success. In addition, big data methods allow correcting for possible interactions and usually result in relatively generalizable findings. To date, however, big data investigations encompassing a large number of participants are still rare in the field of real-time fMRI neurofeedback. The existing ones have either descriptively summarized the field (Heunis et al., 2020; Thibault et al., 2018), or investigated the influence of pre-training brain activation levels on neurofeedback success (Haugg et al., 2020). Here, for the first time, we employ machine learning methods to compute the influence of a wide range of different subject- and study-specific factors on real-time fMRI neurofeedback success. In particular, we investigated the influence of 20 different factors on neurofeedback success in 608 participants undergoing neurofeedback training across 28 independent studies. The investigated factors included three subject-specific factors, six region of interest (ROI)-based factors, and eleven paradigm-specific factors.

Identifying factors that influence neurofeedback success can help to design more effective neurofeedback studies in the future. This can improve neurofeedback studies investigating healthy participants and, more importantly, it can, further, improve clinical neurofeedback interventions. Future designs with increased effectiveness will allow participants to train their target brain regions more efficiently, thus reducing cognitive exhaustion and overall costs. Critically, increasing the effectiveness of neurofeedback designs is an important step towards the alleviation of clinical symptoms, by enabling the development of advanced, personalized treatments for psychiatric and neurological disorders. Taken together, our research aim is to utilize big data approaches in an effort to guide future empirical investigations that utilize realtime fMRI neurofeedback.

## Material and methods

### Included studies

Data for this mega-analysis could not be gathered from publications alone as single subject data were needed. Therefore, we contacted corresponding authors from real-time fMRI neurofeedback studies via i) the mailing list of the real-time functional neuroimaging community (https://utlists.utexas.edu/sympa/subscribe/rtfin), ii) neuroimaging conferences, and iii) direct email communication, in order to ask for data contributions. To ensure generalizability and to generate a dataset sufficiently large for machine learning analyses, we included fMRI-based neurofeedback studies of any training type (activity- as well as connectivity-based neurofeedback), any target brain region(s), and any participant populations. We received data contributions from authors of 28 independent studies (Auer, Schweizer, & Frahm, 2015; Emmert, Kopel, et al., 2017; Hellrung et al., 2018; Kim et al., 2015; Kirschner et al., 2018; Kohl et al., 2019; MacInnes, Dickerson, Chen, & Adcock, 2016; Marins et al., 2015; Marxen et al., 2016; McDonald et al., 2017; Megumi, Yamashita, Kawato, & Imamizu, 2015; Nicholson et al., 2017; Pamplona et al., 2020; Papoutsi et al., 2020, 2018; Scharnowski et al., 2012, 2015; Sorger, Kamp, Weiskopf, Peters, & Goebel, 2018; Spetter et al., 2017; Yao et al., 2016; Young et al., 2017; Zich et al., 2020), covering a wide range of trained brain regions, different study designs, and participant populations. Table 1 provides an overview of all studies that contributed data to this mega-analysis. In total, we collected data from 608 participants, including 229 patients and 379 healthy participants.

**Table 1:** Overview of studies included in the mega-analysis. We received data from 28 independent neurofeedback studies, including 608 participants (229 patients and 379 healthy participants). 24 studies used activity-based neurofeedback, 6 studies used connectivity-based neurofeedback. Abbreviations: ACC – Anterior Cingulate Cortex, dlPFC – dorsolateral Prefrontal Cortex, mPFC – medial Prefrontal Cortex, M1 – Primary Motor Cortex, OFC – Orbitofrontal Cortex, PCC – Posterior Cingulate Cortex, PMC – Pre-Motor Cortex, PHC – Parahippocampal Cortex, SMA – Supplementary Motor Cortex, SMC – Somatomotor Cortex, SPL – Superior Parietal Lobe, VTA – Ventral Tegmental Area.

| **Author (year)** | **ROI(s)** | **participants** | **neurofeedback type** |
| --- | --- | --- | --- |
| Auer et al. (2015) | SMC | healthy (N=16) | activity |
| Emmert et al. (2017) | auditory cortex | tinnitus (N=11) | activity |
| Hellrung et al. (2018) | amygdala | healthy (N=34) | activity |
| Hellrung et al. (in prep) | amygdala | healthy (N=16) | activity |
| Hellrung et al. (in prep) | insula | healthy (N=11) | activity |
| Keynan et al. (in prep) | amygdala | healthy (N=33) | activity |
| Kim et al. (2015) | ACC, mPFC, OFC, PCC, precuneus | tobacco use disorder (N=14) | connectivity, activity |
| Kirschner et al. (2018) | VTA | healthy (N=27), cocaine use disorder (N=24) | activity |
| Kirschner et al. (in prep) | VTA | schizophrenia (N=14) | activity |
| Kohl et al. (2019) | dIPFC | overweight (N=16) | activity |
| Kohl (pilot data) | dIPFC | overweight (N=9) | activity |
| Liew et al. (in prep) | left PMC, left SMA | healthy (N=10) | connectivity |
| MacInnes et al. (2016) | VTA | healthy (N=19) | activity |
| Marins et al. (2015) | left PMC | healthy (N=14) | activity |
| Marxen et al. (2016) | amygdala | healthy (N=32) | activity |
| McDonald et al. (2017) | default mode network | healthy (N=68), psychiatric disorders (N=72) | activity |
| Megumi et al. (2015) | left lateral parietal, left M1 | healthy (N=12) | connectivity |
| Nicholson et al. (2017) | amygdala | PTSD (N=14) | activity |
| Pamplona et al. (2020) | default mode network, sustained attention network | healthy (N=15) | activity |
| Papoutsi et al. (2018) | SMA | Huntington's disease (N=10) | activity |
| Papoutsi et al. (2020) | SMA, left striatum | Huntington's disease (N=16) | connectivity, activity |
| Scharnowski et al. (2015) | SMA, PHC | healthy (N=7) | activity |
| Scharnowski et al. (2012) | visual cortex | healthy (N=10) | activity |
| Sorger et al. (2017) | individually different | healthy (N=10) | activity |
| Spetter et al. (2017) | dIPFC, vmPFC | obesity (N=8) | connectivity |
| Yao et al. (2016) | anterior insula | healthy (N=18) | activity |
| Young et al. (2017) | amygdala | depression (N=18) | activity |
| Zich et al. (2020) | amygdala, dIPFC | adolescents (N=27) | connectivity |

### Neurofeedback success measures

To assess neurofeedback success, we asked authors to provide the average feedback value for each neurofeedback training run. Feedback values were defined as the measures that determined the feedback given to the participants during neurofeedback training. Consequently, the type of feedback values varied between different neurofeedback studies (e.g. percent signal change values, beta values, Bayes factors, correlations values etc.). Based on these feedback values, we then defined two general measures for neurofeedback success that would allow for comparisons between participants of different studies and, more importantly, for pooling all participants together:

- *Neurofeedback performance:* General neurofeedback performance for each participant was calculated based on the ratio of successful neurofeedback training runs as compared to unsuccessful neurofeedback training runs. Successful neurofeedback training runs were defined as runs showing feedback values with positive signs for up-regulation and negative signs for down-regulation. For the classification analyses, participants who showed more than 50% of successful neurofeedback training runs were labelled as successful, the others as unsuccessful.
- *Neurofeedback improvement:* Neurofeedback improvement of each participant was calculated based on the slope of the neurofeedback learning curve, i.e. the slope of the regression line over the feedback values of all neurofeedback training runs. For classification analyses, successful participants were then defined as participants with a slope greater than 0, non-successful participants showed a slope smaller or equal 0.

### Investigated factors influencing neurofeedback performance and neurofeedback improvement

We investigated the influence of 20 different factors on neurofeedback success. These continuous and categorical factors included:

- Three subject-specific factors: (1) age of the participant in years, (2) sex of the participant, (3) health status of the participant (healthy participant or patient);
- Six region of interest (ROI)-based factors: (1) ROI(s) is/are cortical or subcortical, (2) ROI(s) is/are a sensory brain region, (3) ROI(s) is/are part of the default mode network (DMN), (4) ROI(s) is/are part of the salience network, (5) ROI(s) is/are part of the motor network, (6) ROI(s) consist(s) of one brain region or more brain regions;
- Eleven experimental design-specific factors: (1) use of connectivity-vs activity-based measure for feedback computation, (2) use of continuous vs intermittent feedback presentation, (3) use vs no use of functional localizer for defining the trained ROI(s), (4) up- vs down-regulation, (5) use of precise strategy suggestions vs no or open strategy suggestions, (6) use of external (monetary) reward vs no external reward given, (7) use of pre-training no-feedback run (functional runs prior to NFB training, where participants are already asked to modulate their brain signals, however, no feedback over regulation performance is provided) vs no pre-training no-feedback run, (8) length of a single neurofeedback training run in seconds, (9) length of a single neurofeedback regulation block in seconds), (10) number of performed neurofeedback training runs, (11) neurofeedback training on one day vs across several days

### Multivariable predictions of neurofeedback performance and neurofeedback improvement

Individual machine learning analyses were performed in Python (v3.8.3) to identify factors that predict participant-specific neurofeedback performance as well as neurofeedback improvement, using multivariable classification models. For the machine learning models, an *Extra Trees* (ExtraTreesClassifier, scikit-learn library v0.23.1; Pedregosa et al., 2011) approach was used, which is a computationally efficient non-linear classification method. *Extra Trees* implements an ensemble of *Extremely randomized trees* (Geurts, Ernst, & Wehenkel, 2006). Ensemble methods improve the performance of base predictors, e.g. decision trees, by accumulating the predictions of the base predictors via, e.g., majority voting. To obtain diverse predictions from the same base predictors processes that introduce randomness are applied when building the base predictors.

The model performance – the prediction accuracy – was estimated using a nested crossvalidation (CV) procedure (Cawley & Talbot, 2010). In the main CV loop, a shuffle-split data partitioning with 10% of the studies in the testing-set was repeated 100 times, resulting in 100 Extra Trees models (300 trees per model). Feature scaling (z-scoring) and hyper-parameter tuning was carried out within the main CV loop, using the training-data of the current CV loop only. Hyper-parameter tuning was implemented in an inner (nested) CV procedure, so a separate CV was carried out for each repetition of the outer CV loop. The inner CV loops used, again, a shuffle-split partitioning scheme with 10% of the studies in the inner testing set and 50 repetitions. To control model complexity, we restricted the maximum number of possible interactions of a decision tree in the *Extra Trees* ensembles by controlling the number of maximum leaf nodes per tree. The candidate maximum number of leave nodes was randomly drawn between 2 and 32 (50 random draws, RandomizedSearchCV, scikit-learn, v0.23.1). The maximum number of leave nodes that led to the lowest squared error was subsequently used in the outer CV loop.

After hyper-parameter tuning, an Extra Trees model was trained in the main (outer) CV loop using the obtained hyper-parameter and 300 trees with no maximum features. Further, minimum samples split was set to 2, minimum samples leaf to 1, and minimum weight fraction leaf to 0.0. No maximum depth and no maximum samples were chosen, minimum impurity decrease was 0.0, ccp alpha was 0.0, and the class weight was computed from training data.

The obtained model was then tested on the respective hold-out set of the main CV loop. The hold-out set (10% of the studies) was explicitly not used in the inner CV loop. In each repetition of the main CV loop, model prediction accuracy was computed. To counter unbalanced classes (more samples in one class than in the other) weighted accuracy was used (Hastie, Tibshirani, & Friedman, 2001). For that purpose an additional model was trained and tested on a shuffled version of the data in each CV loop.

After obtaining the results of the 100 repetitions of the outer CV loop, we assessed whether the models performed statistically significantly better than chance level by applying a bootstrap test (100,000 bootstrap samples; Efron, 1979). For this test, the null-hypothesis was that the difference between accuracy and chance level is on average smaller or equal to zero (Table 2).

**Table 2:** Extra Trees prediction accuracy for the neurofeedback performance and the neurofeedback improvement target.

| | Average of weighted accuracy ± standard deviation | Average of weighted accuracy at chance level | p value of smaller or equal chance level |
| --- | --- | --- | --- |
| 1. Predicting neurofeedback performance | 60.3% ± 12.3 | 51.0% | p < 0.001 |
| 2. Predicting neurofeedback improvement | 48.1% ± 9.0 | 48.4% | p = 0.614 |

Further, we analyzed the importance of each factor for the overall model performance. In specific, the factor importance was estimated by summing up contributions per factor, over the decision tree splits. The total importance of a feature was then computed as the normalized importance of that feature averaged over the trees in the ensemble (Hastie et al., 2001). Correlation of features leads to a split of this importance measure among these features (see Figure S1 in Supplementary Material for correlation map). To determine whether a feature’s contribution was statistically significant, we tested that feature’s importance against the feature importance obtained by a model that was trained with the same parameters, but shuffled data. The null-hypothesis tested per feature was that the median difference in feature importance is smaller or equal to zero. The null-hypothesis was tested with a bootstrap test (100,000 bootstrap samples per feature; Efron, 1979). Obtained p-values were Bonferroni-corrected for multiple comparisons.

The entire analysis (computing the models and the contributions of factors) was carried out two times. First, to predict neurofeedback performance and a second time to predict neurofeedback improvement.

## Results

### Neurofeedback success

When investigating neurofeedback performance, we observed that 69.41% of all participants were labelled as successful, meaning that for them, more than 50% of all neurofeedback training runs were successful. Only 9.70% of participants were characterized by 25% or less successful runs. On average, participants presented 72.36% successful neurofeedback runs. For neurofeedback improvement over runs, we observed an average slope of 0.09 across all participants. Here, 59.70% of the participants showed a positive slope and, therefore, were able to improve their neurofeedback performance over time (see Figure 1).

**Figure 1:**
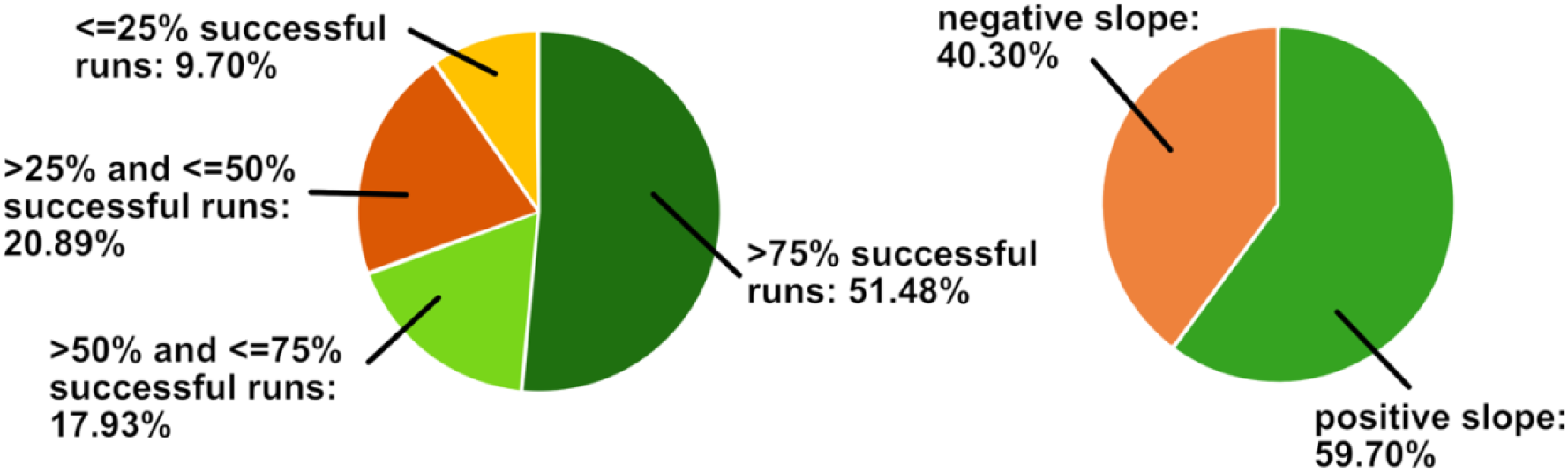
Distribution of neurofeedback success. **Left:** More than half (51.48%) of all participants undergoing neurofeedback training performed 75% or more of successful runs. Only 9.70% of the participants performed 25% successful runs or fewer. **Right:** 59.70% of all participants undergoing neurofeedback training show positive slopes of their learning curves, indicating an improvement over time.

### Prediction accuracy of neurofeedback performance and neurofeedback improvement

The *Extra Trees* machine learning model was able to predict neurofeedback performance from the investigated factors with an average accuracy of 60.3%, which is significantly better than the average accuracy at chance level with 51% (p<.001). However, no prediction better than chance was revealed for neurofeedback improvement (Table 2).

As only the neurofeedback performance measure could be predicted with a better than chance accuracy, only the influence of factors on neurofeedback performance, but not neurofeedback improvement, are valid to be interpreted. Consequently, normalized model-based feature importance was only calculated for the neurofeedback performance target, but not for the neurofeedback improvement target (see Figure 2). Two factors contributed significantly to the prediction result: whether a study included a pre-training no-feedback run (median relative importance 59.3%; Figure 2) and whether a participant was a patient or a healthy participant (median relative importance 31.1%; Figure 2). More specifically, including a pre-training no-feedback run, as well as performing a study with patients increases the chance for a successful neurofeedback run.

**Figure 2:**
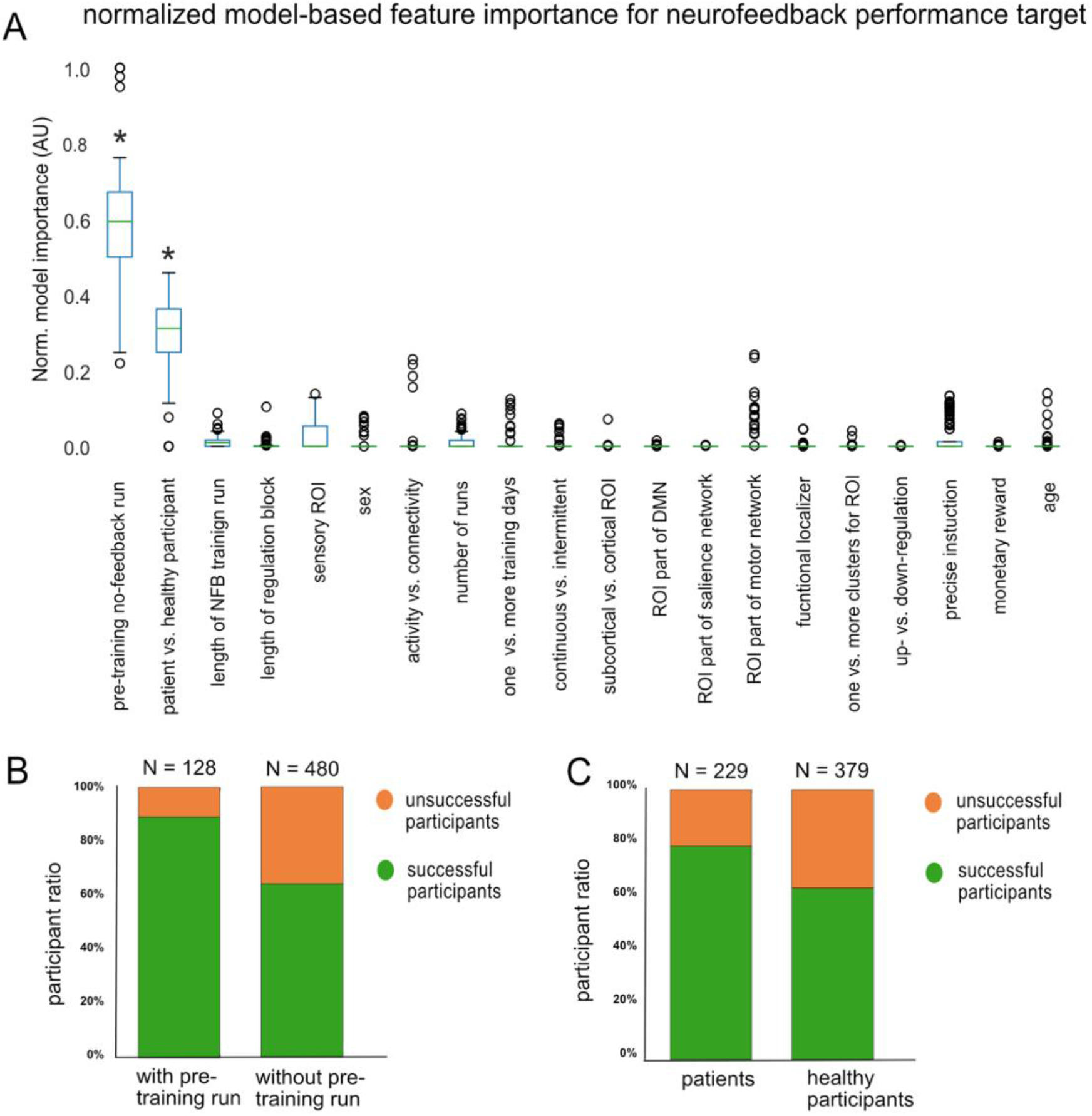
Distribution of feature importance for predicting neurofeedback performance. A Significant normalized model-based feature importance was observed for the feature pre-training no-feedback run and for the feature patient versus healthy participant. B Participants who performed a pre-training no-feedback run were more successful during neurofeedback than participants without a pre-training no-feedback run. C Patients were more successful than healthy participants.

## Discussion

In this study, we investigated the influence of 20 different factors on neurofeedback performance and improvement, including three subject-specific factors, six ROI-based factors, and eleven paradigm-specific factors. When targeting neurofeedback performance, our classification model achieved an accuracy of 60.3%, which was significantly better than chance level. In contrast, classification for the neurofeedback improvement target did not reach an accuracy level above chance level. Overall, we observed considerably high neurofeedback success rates across all 28 studies, with around 60% of all participants showing positive slopes and around 70% of all participants showing more than 50% successful neurofeedback runs. Our results revealed two factors that showed high model importance for the neurofeedback performance classification, suggesting that they may significantly influence neurofeedback performance.

### Factors that influence neurofeedback performance

The first significant factor influencing neurofeedback performance is the presence or absence of a pre-training no-feedback run. Here, significantly higher ratios of successful neurofeedback runs were found for studies that included a pre-training no-feedback run in their study design. Pre-training no-feedback runs are functional runs prior to neurofeedback training, where participants are already asked to modulate their brain signals, however, no feedback regarding regulation performance is provided (e.g. see Kim et al., 2015; Kirschner et al., 2018; MacInnes et al., 2016; Young et al., 2017). These no-feedback runs can serve several purposes, for instance, helping participants familiarize themselves with the neurofeedback paradigm and scanning environment where the following runs will take place. Importantly, they serve as a baseline run for comparisons with subsequent neurofeedback training runs and transfer no-feedback runs after neurofeedback training (Auer et al., 2015; MacInnes et al., 2016). One reason for our finding that pre-training no-feedback runs can benefit neurofeedback performance might be that prior familiarization with the neurofeedback setup and an additional run to practice one’s brain regulation strategies will make it easier for the participants to perform well.

The second factor that demonstrated significant model importance for neurofeedback performance classification was whether a healthy participant or a clinical patient was undergoing neurofeedback training. Specifically, we found that patients showed higher ratios of successful neurofeedback runs than healthy participants. Similar results have already been reported in an empirical neurofeedback study where the authors observed significantly higher default mode network (DMN) upregulation performance in a heterogeneous group of patients, compared to healthy controls (Skouras & Scharnowski, 2019). The authors argued that this finding might be linked to higher observed scores in DMN eigenvector centrality in the patient group than in the control group, i.e. in the patient group the DMN was more strongly connected to the rest of the brain. This is in line with a recent suggestion by Bassett and Khambhati who argue that areas which are strongly functionally connected within the brain (such as it is the case for the DMN) might be easier to be trained with neurofeedback (Bassett & Khambhati, 2017). Further, it is also possible that patients show better performance in neurofeedback regulation due to more dysfunctional brain patterns as compared to healthy subjects, leaving more room for regulation and making ceiling effects less likely. Here, it should be noted that neurofeedback performance might still differ significantly between different patient populations, due to differences in cognitive deficits which might attenuate attention in general and neurofeedback regulation performance in specific (Heeren et al., 2014; Li et al., 2010; Lussier & Stip, 2001). Further, the observed differences in neurofeedback performance between patients and healthy participants might also be driven by differences in the experimental design. Neurofeedback paradigms in clinical populations have oftentimes been piloted more thoroughly, and sometimes even follow a series of several neurofeedback studies in healthy populations which serve as pilots or templates for implementing the optimized final neurofeedback patient studies. For instance, Kirschner et al. (Kirschner et al., 2018) trained participants with cocaine use disorder to regulate their dopaminergic midbrain using a paradigm that had been previously successfully applied to healthy participants (Sulzer et al., 2013). Consequently, high risk studies that are more likely to show a high percentage of unsuccessful neurofeedback runs, e.g. studies using a novel analysis method or an ultra-high-field MRI scanner, might be less often performed with patient populations. Finally, also a difference in the participants’ motivation might influence the better performance of patients as compared to healthy participants. Many patients undergo neurofeedback training in the hope to improve their clinical symptoms while healthy participants mainly participate out of generic interest or in order to receive a monetary compensation. Therefore, it is likely that patients put more effort into the neurofeedback regulation task than healthy participants.

Taken together, our results indicate that it would be beneficial to include a pre-training no-feedback run in order to improve neurofeedback performance. Furthermore, our results demonstrate better neurofeedback performance of patients as compared to healthy participants. While the participant sample is primarily defined by the biological/clinical question under investigation and, thus, does not constitute an open parameter regarding design optimization, this finding nevertheless has strong implications for the design of future neurofeedback studies. Further, our findings emphasize the clinical potential of neurofeedback interventions: Even in cases where only small or moderate effects have been observed in neurofeedback studies on healthy participants, effects in patients might be nonetheless considerably stronger and clinically relevant, based on the same neurofeedback paradigm.

### Features that do not predict neurofeedback performance

Most of the features included in the machine learning analysis did not play an important role with regards to the classification of participants, neither for neurofeedback performance nor neurofeedbackimprovement analyses. One reason for this might be that the majority of our included features were based on parameters specific for each study’s design, such as information on the paradigm or chosen ROI(s), rather than subject-specific features. These design-specific features were deliberately chosen for our analysis to identify parameters that could be easily modified when designing future neurofeedback studies. However, neurofeedback success also varied considerably within single neurofeedback studies (Bray et al., 2007; Chiew et al., 2012; deCharms et al., 2005; Haugg et al., 2020; Johnson et al., 2012; Ramot et al., 2016; F. Robineau et al., 2014; Scharnowski et al., 2012; Yoo et al., 2008), despite all design-specific parameters being identical for the participants of a study. This indicates that subject-specific factors such as biological measures (e.g. heart rate, pulse, stress level), personality traits and cognitive measures, intelligence, the ability to perform mental imagery, or the subject’s attention and motivation (see (Cohen & Staunton, 2019) for a systematic review) might be important factors for successful neurofeedback training. Further, also individual brain-based measures, such as functional connectivity (Scheinost et al., 2014), eigenvector centrality (Skouras & Scharnowski, 2019), or the connectivity of the trained brain region to other higher-order cognitive areas (Bassett & Khambhati, 2017) have been previously discussed as possible factors that might influence neurofeedback success. Due to such information not being available for our data, we were not able to assess the effect that these parameters might have on neurofeedback success. In the future, more harmonization efforts in assessing subject-specific data across differentneurofeedback studies will therefore be necessary.

A complementary reason why many features included in our analysis were not predictive of neurofeedback success was the heterogeneity of the dataset. As we aimed at finding generalizable factors that influence neurofeedback success across a wide range of different neurofeedback studies, we purposely included diverse studies training different ROIs, different participant populations, and using a variety of experimental designs and methods, thus making predictions very difficult. It is possible that by investigating more homogeneous subsets of the data, certain additional factors might become predictive even though they were not predictive when pooling all studies together. However, establishing more homogeneous subsets for solid machine learning analyses will require more data than is currently available.

### Neurofeedback success target measures

Our results were most likely not only driven by the included features, but also by the chosen target measures for neurofeedback success. To date, no commonly accepted measure for neurofeedback success has been established and measures vary between different studies (Haugg et al., 2020; Paret et al., 2019; Thibault et al., 2018). For instance, neurofeedback feedback values during a single neurofeedback regulation block or run can be assessed with a wide variety of different methods, such as percent signal change, beta values, or connectivity values. The heterogeneity of feedback values complicates machine learning approaches that require a common target feature. Even if we had access to the raw imaging data, post-hoc reanalyses with an identical analysis pipeline for all studies would not solve this problem, because such a measure would not reflect the feedback that was provided to the participants during training. Choosing neurofeedback performance and neurofeedback improvement as targets for this mega-analysis allowed for pooling this large set of heterogeneous studies, thus, increasing statistical power and generalizability. In addition, by using a dichotomous classification approach (e.g. positive vs. negative slope), we could, further, account for some of the heterogeneity of our data. For instance, when the slope of a neurofeedback learning curve is computed based on only two runs, the resulting values are more likely to be actual outliers, as compared to when the slope of a neurofeedback learning curve based on 20 runs is calculated (Kwak & Kim, 2017). We avoided this problem by using a classification-based instead of a regression-based machine learning approach.

Furthermore improvement regarding the heterogeneity of the neurofeedback success measures might be expected from developing and establishing a commonly accepted model of neurofeedback learning. To date, the underlying mechanisms of neurofeedback have not been fully determined (Cohen & Staunton, 2019; Sitaram et al., 2016), making it difficult to identify the most important attributes of neurofeedback learning, towards creating a comprehensive neurofeedback success measure. With more neurofeedback data becoming publicly available thanks to the Open Science initiative, another solution might be to only consider studies that used exactly the same feedback success measure while still finding enough data to carry out similar analyses.

## Conclusion

With 59.70% of all participants showing positive slopes and 69.41% of all participants having more than 50% of successful neurofeedback runs, our data indicate that neurofeedback training is overall successful, although with large room for improvement. Using machine learning on the largest neurofeedback data set to date, we were able to identify two measures that might influence neurofeedback success and, thus, could lead to improvements in the efficacy of neurofeedback interventions: Participants who performed a pre-training no-feedback run prior to neurofeedback training and participants who were patients generally performed better. Nevertheless, the medium overall predictability of our analyses indicates that further studies based on larger datasets and including more features are needed. In the future, our mega-analysis machine learning approach combined with increased data availability from homogeneous studies might allow for identifying more crucial factors, designing more efficient neurofeedback studies, improving clinical neurofeedback-based interventions, and understanding better how learning with neurofeedback is accomplished.

